# Filoviruses Can Efficiently Infect Human Neuron-Like Cells Without Genetic Adaptation

**DOI:** 10.1101/2019.12.12.874016

**Authors:** Alexander J. McAuley, Mary Tachedjian, Glenn A. Marsh

## Abstract

Recent large-scale Ebola outbreaks, combined with improved follow-up of survivors, has permitted the observation of common long-term neurological sequelae in patients that have survived Ebola virus infection. To date there have been few studies into neurological infections by Ebola or related filoviruses, however, recent studies have isolated infectious virus from patients’ cerebrospinal fluid months after being discharged from the treatment facility.

In order to determine whether different filoviruses were capable of infecting human neurons, the human neuroblastoma cell lines, SH-SY5Y and M17, were chemically-differentiated into more neuron-like cells using established protocols. The neuron-like profiles of the differentiated cells were confirmed by the determination of expression of a range of neuron-specific markers. Zaire ebolavirus, Reston ebolavirus, and Marburg virus were serially-passaged in both cell lines to determine permissiveness of the cells, as well as permit the acquisition of adaptive mutations in the viral genomes. Whilst Marburg virus grew to high titres in both cell lines, Zaire ebolavirus only grew in SH-SY5Y cells, and Reston ebolavirus rapidly died out in both cell lines. Whole-genome sequencing of the passaged viruses revealed two consensus-level non-coding mutations in the SH-SY5Y-passaged Marburg virus. Viral growth kinetics were determined for pre- and post-passaging Zaire ebolavirus and Marburg virus in both human neuronal cell lines, as well as the human hepatocyte cell line, Huh7. Growth kinetics were similar for both the pre- and post-passaged viruses, suggesting that adaptive mutations were not required for efficient growth in these cells.

This study is the first to demonstrate that filoviruses are capable of infecting human neuron-like cells in a species-specific manner. Marburg virus-infected cells remained alive up to Day 21 post-infection, suggesting that long-term neurological sequelae following filovirus infection may be a result of direct neuronal infection, and that infection of neurons might contribute to viral persistence in survivors.

**Author Summary:** Filoviruses, including Ebola and Marburg viruses, have been traditionally considered “haemorrhagic fever” viruses, with infections causing bleeding and frequently death. Recent large-scale outbreaks in Africa have challenged these assumptions due to a significant number of patients reporting neurological symptoms sometimes months after infection. In many of these patients, virus was present at detectable levels in the fluid surrounding the brain. There has been significant debate about the ability of Ebola and Marburg viruses to infect and grow in human neurons (brain cells), and evidence has been lacking due to the lack of feasibility in taking brain samples. Our study demonstrates that both Zaire ebolavirus and Marburg virus are capable of infecting cells derived from human brains without needing to change, and without killing the cells. Reston ebolavirus, a related virus that appears not to cause disease in humans, was not able to grow efficiently in these cells. Our findings show that these viruses might be capable of living in the brains of survivors for long periods of time, similar to previous observations in the eye and testes. In addition, the response of the body to these infected cells might account for the neurological symptoms described by patients.

## Introduction

The family *Filoviridae* contains some of the most virulent human pathogens known. Members of this family, which includes the Ebola and Marburg viruses, are associated with mortality rates in excess of 50% (1). A recent large-scale outbreak of Zaire ebolavirus in western Africa has resulted in tens of thousands of infections, with over 10,000 fatalities, whilst an ongoing outbreak in the Democratic Republic of the Congo has claimed over 2,000 lives with over 3,000 confirmed cases (1, 2). For a long time, filoviral infections were considered “haemorrhagic fevers”, affecting peripheral organs of the body including the liver, spleen, and lungs (3–5). Little attention was focussed on neurological involvement during filovirus infection, despite the observation of Marburg virus-associated encephalitis during the first recorded filovirus outbreak in 1967 (6). Following this outbreak, a study demonstrated panencephalitis in mice inoculated intracerebrally with a mouse-adapted Marburg strain (7)

Subsequent to the initial Marburg virus outbreaks, most symptomatic human filovirus infections occurred in remote communities in Africa, in countries without efficient medical infrastructure and a small number of survivors with minimal opportunities for follow-up. A recent large-scale Ebola virus outbreak in West Africa has permitted the tracing of survivors and have allowed for the determination of longer-term effects of Ebola virus infection in humans (8–20). These long-term sequelae include a range of neurological and psychological disorders, including headache, fatigue, depression, meningoencephalitis, and cerebral and/or cerebella atrophy (8–10, 12–17, 19, 20).

Due to the risk and complexities associated with the collection of anatomical samples from Ebola patients, analysis of brain material from infected individuals has not been possible. However, a number of studies have demonstrated the presence of infectious virus within the cerebrospinal fluid (CSF) of Ebola survivors, months after initial discharge from hospital (8, 15, 21). In addition, there have been numerous reports of ocular involvement following Ebola virus disease, leading to persistent infection (22–26). Taken together, these observations demonstrate that Ebola virus is capable of crossing both the blood-ocular and blood-brain barriers to gain entry to otherwise immune-privileged sites.

The exact mechanisms responsible for the neurological signs and symptoms observed during and following Ebola and Marburg infections are poorly understood but are likely to involve direct infection of human neurons and/or immunopathology due to infection-associated immune responses. Direct infection of neurons by Ebola and Marburg viruses remains a controversial topic, due to the lack of direct experimental evidence, although some groups have suggested that infection of the optic nerve could account for the subsequent generation of retinal lesions (26).

One of the key reasons for the lack of evidence for neural infection is that there are no useful animal models of persistent filovirus infection. Within human patients, it is common for neurological signs to present months after surviving the initial infection. Animal studies of filovirus infection tend to be lethal, with experimental endpoints of just a few days or weeks. A recent retrospective study of historical samples from surviving non-human primate Ebola infections aimed to characterise persistent, subclinical virus infection. However this study was limited by the fact that samples were only available up to 43 days post-infection, and most of the surviving animals were either vaccinated or had received experimental treatments (27).

Some *in vitro* studies using vesicular stomatitis virus (VSV)-based pseudoviruses with Ebola virus glycoproteins present on the surface of the virion have demonstrated that these pseudoviruses are capable of infecting neurons in neonatal mice, leading to neurodegeneration and death in these animals (28), although the same observations were not made in adult mice or non-human primates (28, 29).

Human neuroblastoma cell lines, including SH-SY5Y and BE(2)-M17 cells, are commonly used for *in vitro* analysis in neuroscience research into conditions such as Alzheimer’s and Parkinson’s disease, and are widely considered to be appropriate *in vitro* models for human neurons given their expression of neuron-specific factors and markers (30–35). The quality of the neuron-like phenotypes (both morphological and biochemical) of these cell lines can be increased through the use of chemical differentiation, often involving drugs such as *trans*-retinoic acid (RA) or staurosporine (36–38). A recent study by Filograna et al (2015) demonstrated that, from the standpoint of a neuronal phenotype, differentiation of M17 cells is better with RA, whereas SH-SY5Y cells differentiate better with staurosporine (37). The resulting differentiated neuron-like M17 cells have been shown to possess a more dopaminergic phenotype, whilst staurosporine-differentiated SH-SY5Y cells possess a more adrenergic phenotype (37). These cells can therefore be used as models for two clades of neurons found within the brain.

Both differentiated and undifferentiated SH-SY5Y and BE(2)-M17 cells have been previously used for *in vitro* studies with a range of neurotropic viruses, including herpes viruses (39–41), neurotropic enteroviruses (42, 43), chikungunya virus (44), poliovirus (45), various orthobunyaviruses (46), henipaviruses (47, 48), and Zika virus (49). In a number of these studies, growth in SH-SY5Y and BE(2)-M17 cells differed based upon whether the strains involved were virulent or attenuated *in vivo*, with neurovirulent viruses reaching higher titres than attenuated ones (43, 45).

Given the uncertainty surrounding the infection of human neurons by Ebola and Marburg viruses, and the previously-demonstrated suitability of both SH-SY5Y and BE(2)-M17 cells to model human neurons *in vitro*, the ability of Zaire ebolavirus, Reston ebolavirus, and Marburg virus to infect and replicate within these cells was characterised. The results demonstrated that Marburg virus grew efficiently in both cell lines tested, while infection with Zaire ebolavirus was limited to SH-SY5Y cells. Reston ebolavirus failed to grow efficiently in either human neuron-like cell type, in keeping with its apparent attenuated phenotype in humans.

## Results

### Chemically-Differentiated BE(2)-M17 and SH-SY5Y Cells are Human Neuron-Like and Express Mature Neuron Markers with Increased Neurite Generation

Treatment of BE(2)-M17 and SH-SY5Y cells with RA or Staurosporine, respectively, resulted in the adoption of a neuron-like morphology (as previously described (37)) marked by the formation of inter-connected neurites, particularly with the differentiated SH-SY5Y cells (Figure 1A). Phenotypic changes to the RA-treated BE(2)-M17 cells were not as pronounced as the staurosporine-treated SH-SY5Y cells, but neurite formation was nonetheless greater than in the mock- or DMSO-only-treated cells.

**Figure 1:**
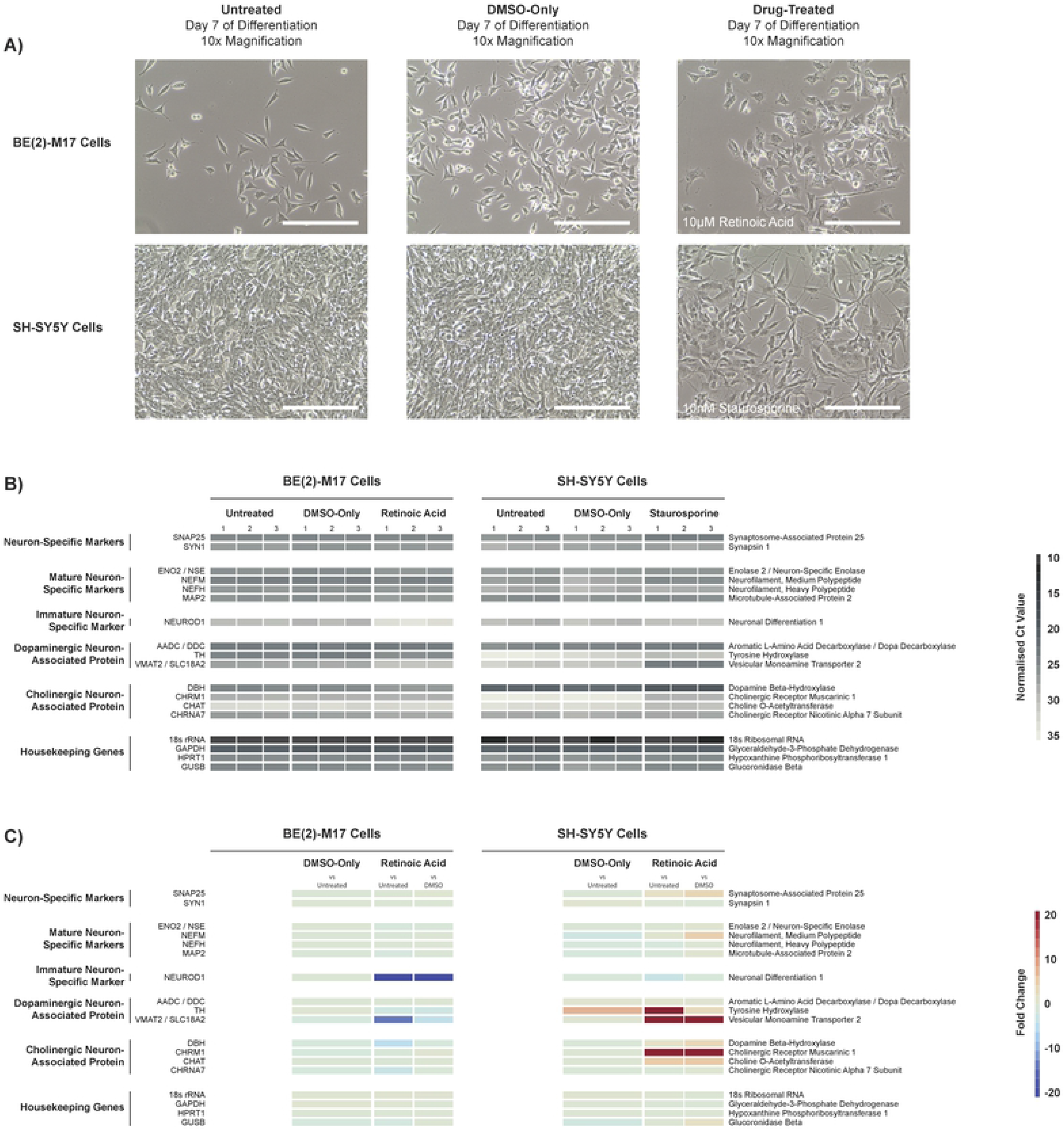
Effect of Chemical Differentiation on Phenotype and Expression Profiles of BE(2)-M17 and SH-SY5Y Cells. BE(2)-M17 and SH-SY5Y cells were either untreated, treated with DMSO-only, or treated with 10μM *trans*-Retinoic Acid (BE(2)-M17 cells) or 10nM Staurosporine (SH-SY5Y cells). Treatments were performed for 7 days as described in the methods section. After the 7-day differentiation, each of the cell condition combinations were imaged to determine cell morphology (A). Total RNA was purified from each cell condition for expression analysis using custom-designed RT-qPCR arrays to determine expression of neuron-specific markers. Ct values were normalised and scaled to generate a pseudonorthern blot of relative RNA levels (B). Fold changes between DMSO-only treatment and untreated cells, as well as drug treatment compared to both DMSO-only treatment and untreated cells, were determined using the ΔΔCt method (C).

In order to determine the suitability of BE(2)-M17 and SH-SY5Y cells as an *in vitro* model of human neurons, expression of commonly-used markers of neuron identity and maturity was determined through the use of custom qPCR arrays. Ct values were normalised to the geometric mean Ct values of the housekeeping genes. The normalised Ct values were scaled and plotted as a semi-quantitative pseudo-Northern Blot to allow for gene expression to be visualised (Figure 1B). Fold change calculations were performed for each cell type using the ΔΔCt method comparing the DMSO-only samples with the equivalent untreated samples, and the drug-treated samples with both the DMSO-only and untreated samples (Figure 1C).

Untreated, mock-treated, and differentiated BE(2)-M17 and SH-SY5Y cells had substantial expression of the general neuronal cell markers SNAP25 and SYN1 (normalised Ct values between 22 and 26). Similarly, expression of the mature neuron specific markers ENO2/NSE, NEFM, NEFH, and MAP2 was high (Normalised Ct values between 22 and 26). The expression levels of these factors were similar in both human neuroblastoma cell lines, and were minimally altered by chemical differentiation, suggesting a fixed, mature human neuron profile. This observation was further supported by the fact that the expression of the immature neuron marker, NEUROD1, was low (Ct values >29) in both cell types. Unlike the main neuron markers, NEUROD1 expression was altered following differentiation, with a 4.6-fold decrease in expression in differentiated SH-SY5Y cells compared to the untreated SH-SY5Y cells. Within BE(2)-M17 cells, the decrease in NEUROD1 expression was even more substantial, with 33.3-fold lower expression in RA-treated cells compared to the untreated cells.

Chemical differentiation of the BE(2)-M17 and SH-SY5Y cells led to more substantial changes in the expression of factors associated with neuron subtypes. Expression of the dopaminergic neuron-associated factors Aromatic L-Amino Acid Decarboxylase (AADC), Tyrosine Hydroxylase (TH), and Vesicular Monoamine Transporter 2 (VMAT2) was strong in untreated BE(2)-M17 cells, with CT values around 22–24 for AADC and TH, and 26 for VMAT2. RA-treatment had little effect on expression of AADC and TH, but led to a 14.7-fold decrease in VMAT2 expression. Expression of cholinergic neuron-associated factors in BE(2)-M17 cells were generally lower than the dopaminergic-associated factors (Ct values >25), and were moderately-affected by RA-treatment (up to 5.9-fold decrease in expression).

In contrast to the BE(2)-M17 cells, differentiation of SH-SY5Y cells generally led to an increased in expression of neuron subtype-associated factors. Treatment with staurosporine led to 35.9- and 155-fold increases in gene expression of the dopaminergic neuron-associated factors TH and VMAT2 in SH-SY5Y cells compared to the untreated cells. Unlike the RA-treated BE(2)-M17 cells, staurosporine treatment of SH-SY5Y cells also led to the increased expression of cholinergic neuron-associated factors DBH (2.5-fold increase compared to untreated), CHRM1 (61.0-fold increase), and CHAT (6.2-fold increase).

### BE(2)-M17 and SH-SY5Y Cells Moderately-Express Known Filovirus Receptors

In addition to characterising neuron-associated factors within the BE(2)-M17 and SH-SY5Y cells, expression of known and potential filovirus receptors was determined by qPCR. As for the neuronal markers, the normalised Ct values were scaled and plotted as a semi-quantitative pseudo-Northern Blot to allow for gene expression to be visualised (Figure 2A), and fold change calculations were performed (Figure 2B). In general, many filovirus receptors do not appear to be expressed strongly in human neuron-like cells. The receptors TIM-1, FCN1, CD209/DC-SIGN, and CLEC4M/L-SIGN each have Ct values >35. Similarly, FOLR1 is expressed with a Ct value >30.

**Figure 2:**
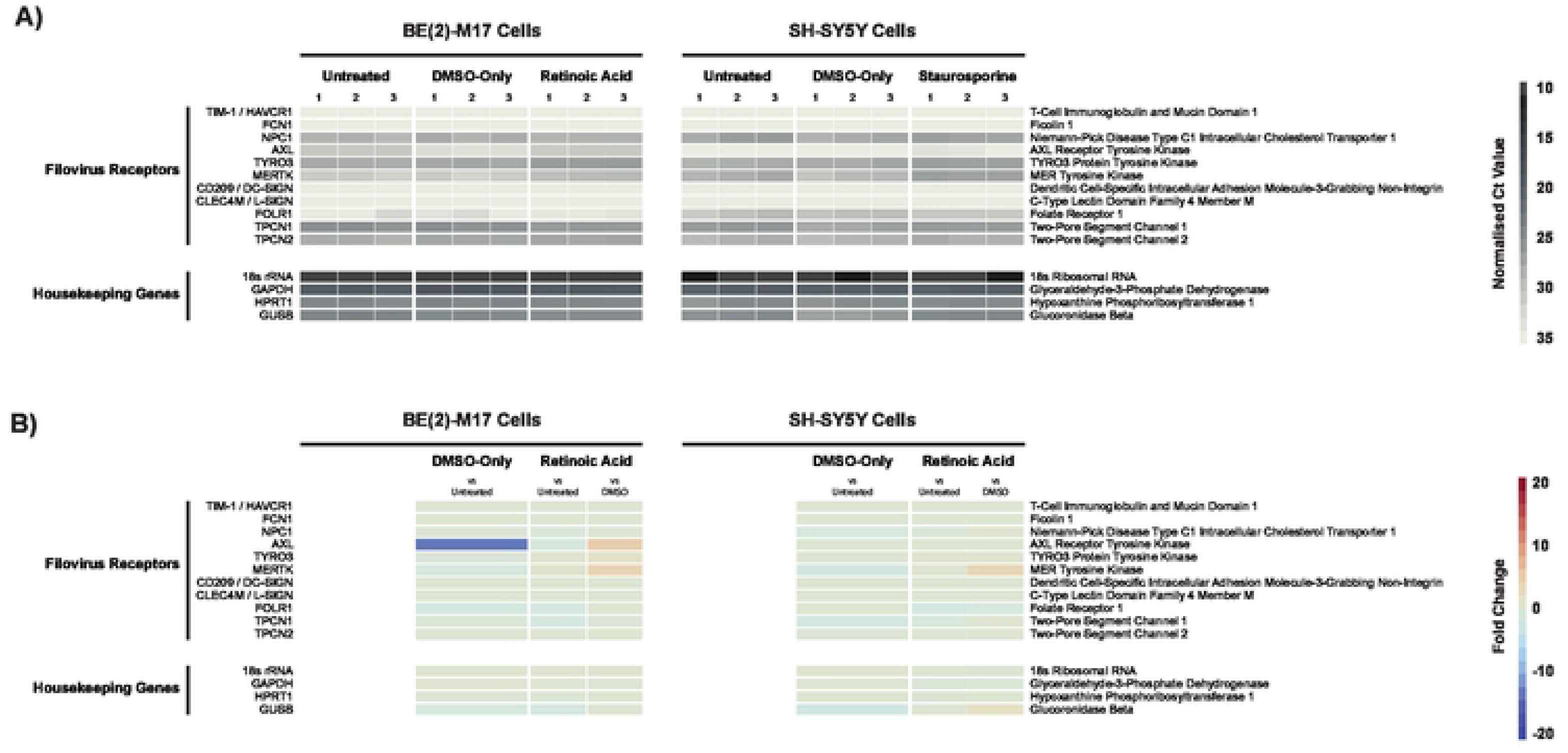
Expression of Known Filovirus Receptors in BE(2)-M17 and SH-SY5Y Cells. BE(2)-M17 and SH-SY5Y cells were either untreated, treated with DMSO-only, or treated with 10μM *trans*-Retinoic Acid (BE(2)-M17 cells) or 10nM Staurosporine (SH-SY5Y cells). Treatments were performed for 7 days as described in the methods section. After the 7-day differentiation, total RNA was purified from each cell condition for expression analysis using custom-designed RT-qPCR arrays to determine expression of known filovirus receptors. Ct values were normalised and scaled to generate a pseudonorthern blot of relative RNA levels (B). Fold changes between DMSO-only treatment and untreated cells, as well as drug treatment compared to both DMSO-only treatment and untreated cells, were determined using the ΔΔCt method (C).

The main surface receptors expressed in the two neuronal cell types are the TYRO3 family of receptor tyrosine kinases (AXL, TYRO3, and MERTK) and the two-pore segment channel proteins (TPCN1 and TPCN2). Within RA-differentiated BE(2)-M17 cells, AXL, TYRO3, and MERTK had Ct values of 30, 26, and 29, respectively. Within the staurosporine-differentiated SH-SY5Y cells, these genes had Ct values of 33, 26, and 26, respectively. The two-pore segment channel proteins, TPCN1 and TPNC2 had Ct values of 25 and 26, respectively, in BE(2)-M17 cells, and 25 and 27 in SH-SY5Y cells.

The only filovirus receptor known to be necessary for viral entry is the intracellular receptors Niemann-Pick C1 (NPC1) protein. Expression of this gene was detected in both cell lines, with Ct values of 28 and 26 in differentiated BE(2)-M17 and SH-SY5Y cells, respectively. Chemical differentiation did not substantially affect expression of NPC1 in either cell type (<1.5-fold change).

### Human Neuron-Like Cells Support Infection and Replication of Some, But Not All, Filoviruses

As a result of their strong neuron-like properties, chemically-differentiated BE(2)-M17 and SH-SY5Y cells were infected with EBOV, RESTV, and MARV at an MOI of 0.5 (based upon virus titres in VeroE6 cells) to determine their permissiveness for filovirus infection. Supernatant samples were harvested at Day 5 post-infection and were used to infect freshly-differentiated cells. The viruses were serially passaged five times to determine whether productive infections could be maintained in human neurons. Supernatant samples collected at each passage were titrated to determine the growth abilities of each virus in the two cell lines (Figure 3).

**Figure 3:**
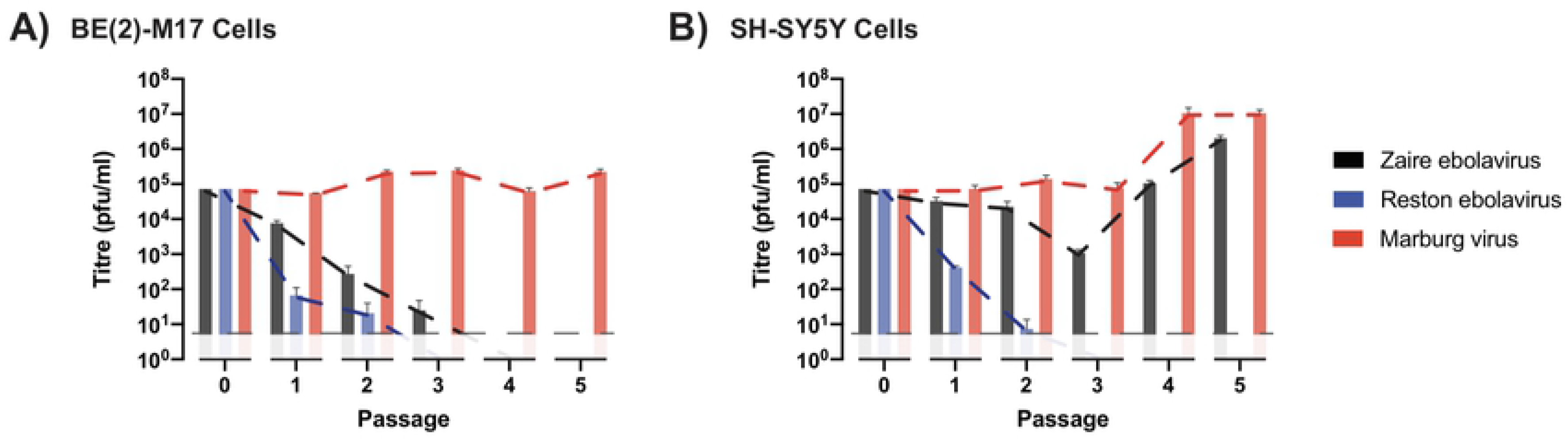
Viral Titres Following Serial Passage of EBOV, RESTV, and MARV in Differentiated BE(2)-M17 and SH-SY5Y Cells. EBOV, RESTV, and MARV were serially passaged five times in triplicate in differentiated BE(2)-M17 and SH-SY5Y cells starting with an MOI of 0.5. Viral titres were determined for each passage. Values plotted are mean titres with standard error.

Growth of EBOV was restricted in BE(2)-M17 cells, with output titres lower than input titres over the first three passages until they dropped below the limit of detection at Passage 4. By contrast, within SH-SY5Y cells, EBOV grew to consistently high titres (>10^5^ pfu/ml). RESTV was unable to be detected by Passage 3 in both cell lines, suggesting that this virus is incapable of productive infection of human neurons. Unlike EBOV and RESTV, MARV grew well in both neuron-like cell lines with titres >10^5^ pfu/ml in BE(2)-M17 cells and >10^7^ pfu/ml in SH-SY5Y cells.

### Passaging of Filoviruses in Human Neuron-Like Cell Lines Results in Minimal Sequence Changes

In order to determine whether passaging of filoviruses in neuron-like cells resulted in the acquisition of adaptive changes in the viral genome, all of the virus-containing Passage 5 samples, and the stock virus used for the initial infections, were sequenced by Illumina next-generation sequencing. The sequences were *do novo* assembled and aligned to the published sequences for each virus strain (EBOV: JQ352763; MARV: KY047763) (Table 1; Supplementary Table 1).

**Table 1:**
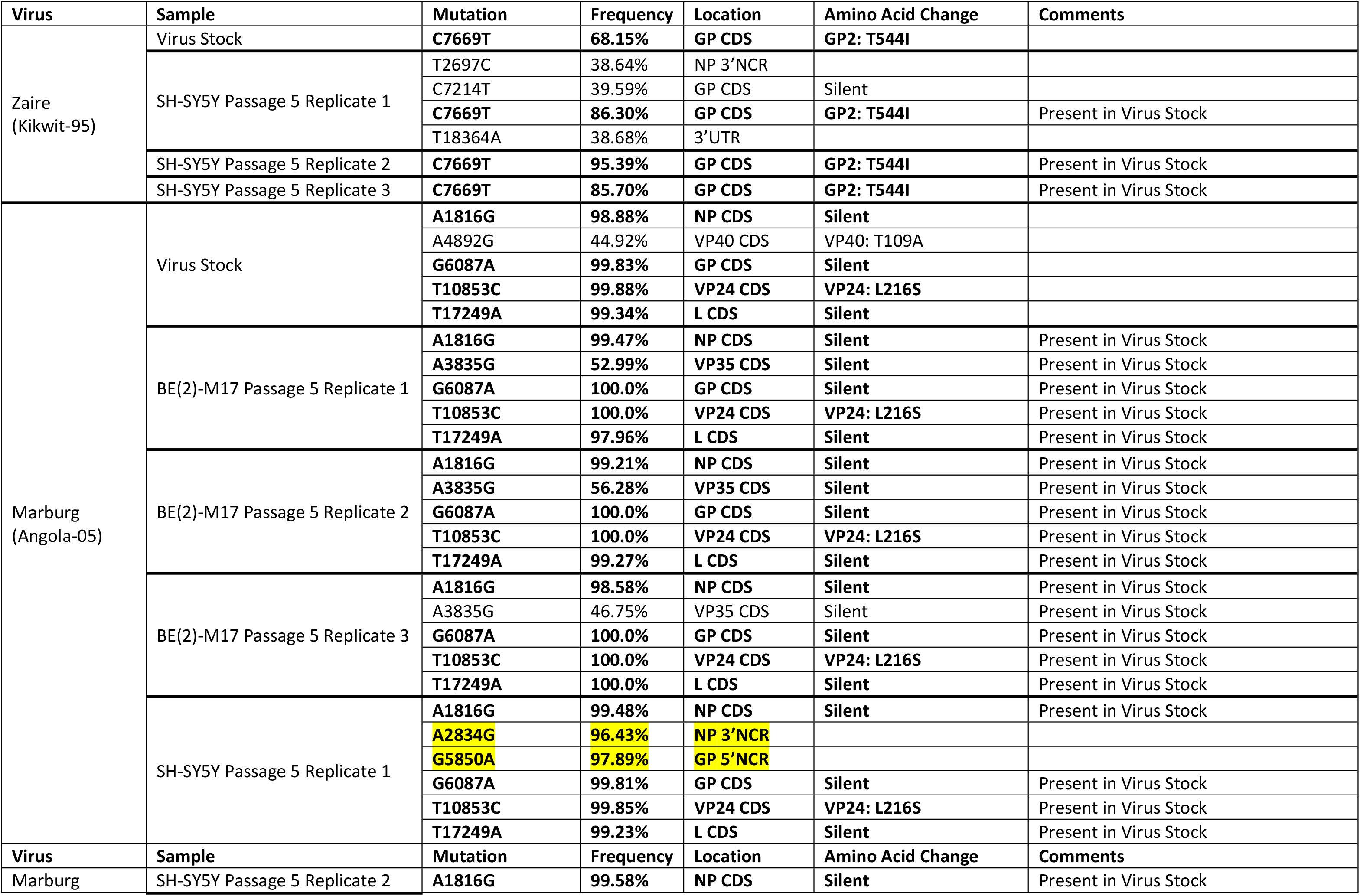
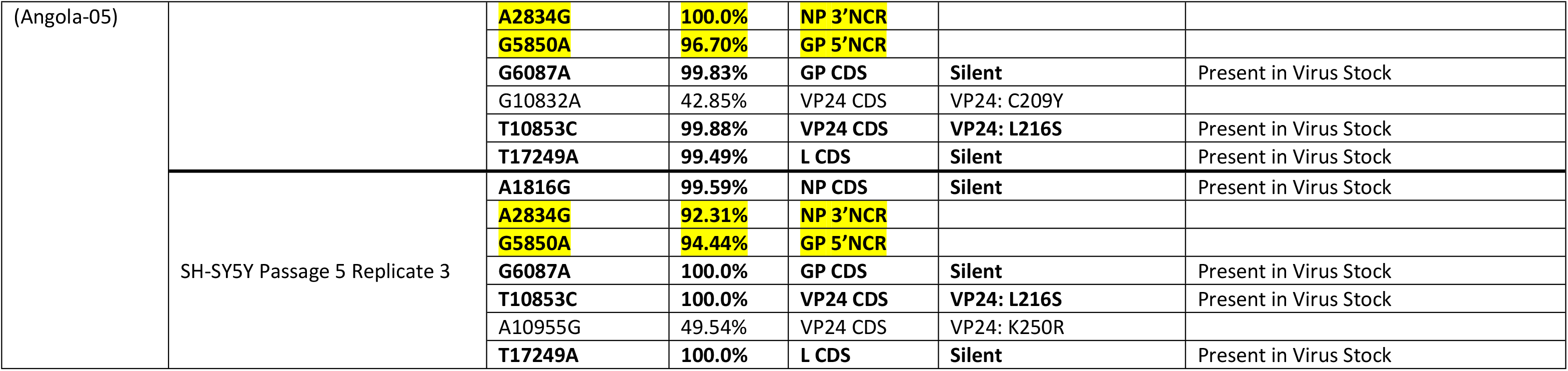
Mutations Present in Passaged EBOV and MARV Samples (>35% Frequency)

The EBOV stock possessed a sub-consensus level (68.2% frequency) T544I amino acid substitution in the GP2 coding sequence compared to the published sequence. Passaging of EBOV in SH-SY5Y cells resulted in an apparent selection for this mutation, with GP2 T554I mutation frequencies >85% in all three replicates of the Passage 5 viruses. In addition to the mutation in GP2, Replicate 1 possessed an additional three minor (<40% frequency) substitutions that were located within non-coding regions.

The MARV stock virus possessed five sequence variations >35% frequency compared to the published sequence for the Angola-05 strain. Four of these mutations were at the consensus level (>90% frequency), and the fifth had a frequency of 44.9%. Of the five mutations, three were silent, and the other two led to semi-conservative amino acid changes in VP40 (T109A; 44.9% frequency) and VP24 (L216S; 99.9% frequency).

None of the differentiated BE(2)-M17 cell-passaged MARV replicates contained additional mutations with >35% frequency compared to the virus stock. However, in Replicates 1 and 2, the T109A substitution in VP40 increased in frequency from 44.9% in the virus stock to 53.0% and 56.3% in the two replicates, respectively. Replicate 3 only saw an increase in frequency to 46.8%.

Passaging MARV in differentiated SH-SY5Y cells resulted in the selection for two consensus-level (>90% frequency) mutations (A2834G and G5850A nucleotide substitutions in cRNA). Neither of these mutations were located within coding sequence, but arose in tandem in all three of the independent replicate virus passage lineages. In addition to these two common non-coding mutations were a VP24 C209Y (42.9% frequency) amino acid substitution in Replicate 2, and a VP24 K250Y (49.5% frequency) amino acid substitution in Replicate 3.

### Efficient Growth of EBOV and MARV in Neuron-Like and Non-Neuron-Like Cells Does Not Require Adaptive Mutations

In order to better characterise the growth of ZEBOV and MARV in human neuron-like cells, and to identify any effects of virus passaging in neuron-like cells on growth in neuron-like and non-neuron-like cells, comparative growth kinetics were performed in Huh7 human hepatocellular carcinoma cells, as well as differentiated BE(2)-M17 and SH-SY5Y cells. Given that EBOV died out in the BE(2)-M17 cells, and MARV passaging in these cells resulted in minimal genome sequence changes, only the stock viruses and the SH-SY5Y Passage 5 viruses (Replicate 3 for EBOV and Replicate 2 for MARV) were used for growth kinetics infections. Cells were infected with a low MOI (0.01) to better tease out any differences in growth (Figure 4).

**Figure 4:**
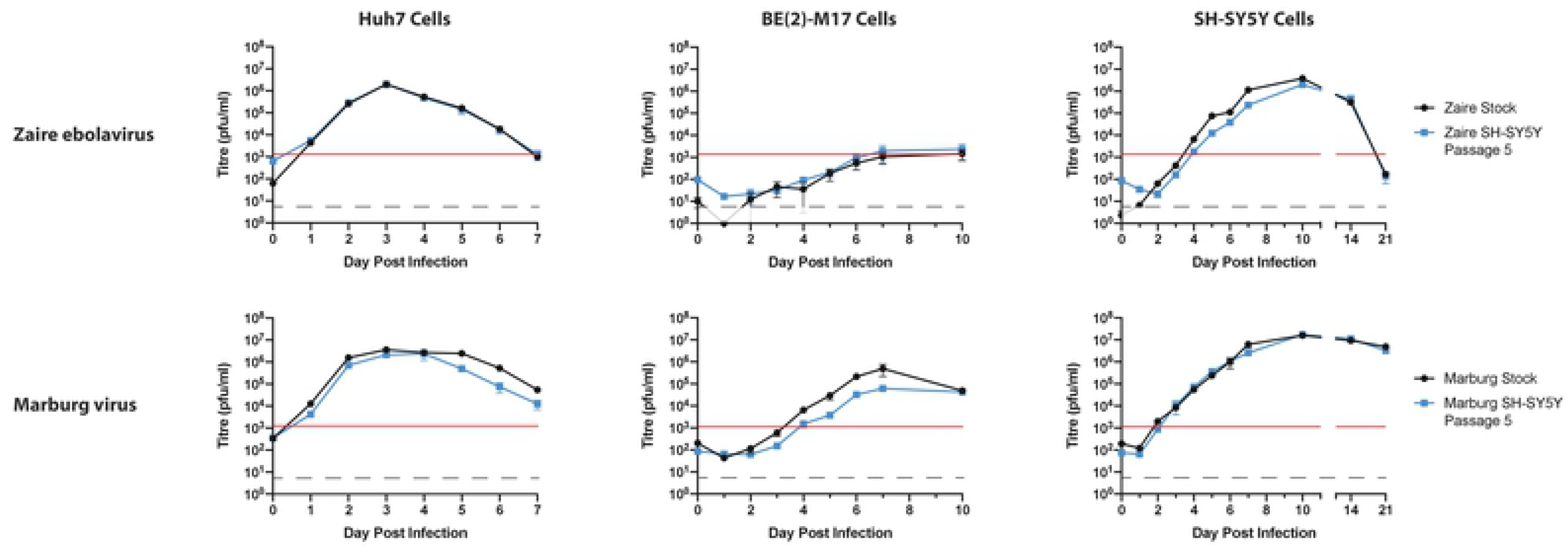
Comparative Growth of Passaged and Stock EBOV and MARV in Huh7 Cells and differentiated BE(2)-M17 Cells. Triplicate flasks of Huh7 cells and differentiated BE(2)-M17 and SH-SY5Y cells were infected with EBOV and MARV from SH-SY5Y Passage 5 or from the original virus stock at an MOI of 0.01. For each of the cell types, daily supernatant samples were harvested on Day 0 to Day 7 of the infection for titration. For the BE(2)-M17 cells, additional samples were taken on Day 10; and for SH-SY5Y cells, additional samples were taken on Day 10, 14, and 21 post-infection. Values plotted are mean titres with standard error.

Growth of EBOV in the three cell lines appeared to be minimally-affected by the viral passaging in SH-SY5Y cells. In all three cell lines, mean titres for the stock and SH-SY5Y-passaged viruses were within 10-fold of one another from Day 1 onward. In Huh7 cells, both the stock and passaged viruses grew to peak titres on Day 3, with a steady decline in titres thereafter corresponding with uniform cell death in the flasks. Interestingly, in BE(2)-M17 cells, EBOV growth titres failed to exceed the amount of virus used in the original infection (red line), which would account for the dying out of the virus during serial passaging in these cells. Growth of EBOV in SH-SY5Y cells was efficient, reaching titres >10^6^ pfu/ml on Day 10 post-infection. Unlike growth in Huh7 cells, EBOV appeared not to readily induce cell death in either the BE(2)-M17 cells or SH-SY5Y cells. This is evidenced by the maintained peak titre to Day 10 post-infection in BE(2)-M17 cells and the high (>10^5^ pfu/ml) titre on Day 14 in SH-SY5Y cells.

Growth of stock MARV compared to SH-SY5Y-passaged MARV showed some differences in Huh7 and BE(2)-M17 cells, but minimal differences in SH-SY5Y cells. In Huh7 cells, both the stock and passaged MARV grew similarly during the first four days, but titres somewhat diverged thereafter, with the titres of passaged MARV declining 24h before those of the stock MARV. In the BE(2)-M17 cells, stock MARV appeared to grow better than SH-SY5Y-passaged MARV from the start, with stock MARV reaching a peak titre on Day 7 almost 10-fold higher than the peak titre for passaged MARV at the same timepoint. Despite this difference in peak titre, both viruses had identical titres on Day 10 post-infection. Unlike the growth in Huh7 and BE(2)-M17 cell, growth of both the stock and passaged MARV in SH-SY5Y cells was almost identical, with high titres maintained to the Day 21 timepoint.

Given the extreme similarity in growth kinetics between the stock and passaged MARV in SH-SY5Y cells, RNA was purified from supernatant samples taken from the triplicate flasks of stock MARV growing in SH-SY5Y between Day 2 to Day 5 of the infection. The samples were sequenced to determine whether the two non-coding mutations identified in the passaged MARV samples arose during growth of stock MARV in SH-SY5Y cells, accounting for the similar grow phenotype. Surprisingly, no sign of the mutations was observed in any of the triplicate samples up to, and including, Day 5 post-infection. This suggested that the mutations were not necessary for efficient growth in SH-SY5Y cells.

## Discussion

The observation of common neurological sequelae in survivors of recent filovirus outbreaks has reignited the debate about whether these observed clinical signs are a direct result of neuronal infection or are a by-product of antiviral immune responses. The demonstration of infectious virus in the CSF of patients is a clear indication that the virus is capable of entering and replicating within the immunoprivileged locale of the CNS (8, 15, 21). The demonstration of viral persistence in other immunoprivileged sites, such as the eye and testes, also raises the question of viral persistence in the CNS (23, 26, 50, 51).

To date there have been few studies looking at the possibility of neuronal infection by filoviruses. This is partly due to the long-standing belief that filoviruses solely affect peripheral organs, and partly due to the lack of demonstrated animal models recapitulating long-term sequelae observed in human survivors. Due to this lack of appropriate animal models, and the dearth of primary human samples, investigation of filovirus neurovirulence is dependent upon the use of *in vitro* tools.

The human neuroblastoma cell lines SH-SY5Y and BE(2)-M17 are two of the best-characterised cell lines for studies into human neurological disease, such as Alzheimer’s and Parkinson’s diseases (30–35), and neurovirology (39–46, 49, 52). Previous studies have confirmed their suitability as *in vitro* models of human neurons due to their expression of human neuron-specific markers, and the fact that their neuron-like phenotypes can be further improved through the use of chemical differentiation agents, such as RA and staurosporine (36–38, 43).

Our gene expression analysis of differentiated BE(2)-M17 and SH-SY5Y cells is in agreement with previous studies into the suitability of these cell lines as *in vitro* models of human neurons, due to the high level of expression of neuron-specific markers (36, 37). Previous studies by Filograna et al (2015) determined that, following RA-differentiation, BE(2)-M17 cells adopted a dopaminergic neuron phenotype, whilst staurosporine treatment of SH-SY5Y cells led to a more cholinergic neuron phenotype, due in part to the expression of factors involved in the conversion of dopamine to noradrelanline (37). More specifically, RA treatment primarily led to a decrease in VMAT2 expression, whilst staurosporine treatment of SH-SY5Y cells led to increased VMAT2 expression, observations in keeping with our gene expression data. This suggests that chemical differentiation of BE(2)-M17 and SH-SY5Y cells leads to the generation of human neuron-like cells of different subtypes. These differences could explain, in part, the differences in permissibility for the filoviruses studied.

The inability of RESTV to grow productively in either of the neuronal cell types is in agreement with the idea that RESTV is not a pathogenic filovirus in humans. Indeed, even in VeroE6 cells, an interferon-deficient cell line in which EBOV and MARV can grow to titres >10^6^ pfu/ml, RESTV will barely grow to 10^4^ pfu/ml (53).

EBOV appears to show interesting neuronal cell-specificity: the virus is capable of growing productively in SH-SY5Y cells, but not in BE(2)-M17 cells. It is clear from the growth kinetics data that replication is occurring in BE(2)-M17 cells, but it is significantly restricted. This would suggest that either the virus is replicating well in only a few cells, or it is replicating poorly in all of the cells. Given the similarity in filovirus surface receptor expression compared to SH-SY5Y cells (determined by normalised Ct value), one explanation could be the difference in expression of NPC1. In BE(2)-M17 cells, the Ct value for NPC1 was only 28 compared to 26 in the SH-SY5Y cells, representing a 4-fold difference in expression levels. It is possible that minimum expression threshold for productive infection with EBOV lies between 26 and 28, which could explain the differential infectability. Another possibility is that there is an as-yet unidentified filovirus receptor differentially expressed in BE(2)-M17 and SH-SY5Y cells.

A recent study by Zapatero-Belinchón *et al* (2019) has suggested that SH-SY5Y cells are a filovirus-resistant cell line due to a lack of filovirus cell surface receptors (54). At first glance it is not clear why this study did not show productive infection with EBOV or MARV in light of the data presented here. One possibility is that the cells they were using were at a high passage, which is anecdotally believed to reduce the neuron-like properties of the cell line and preclude chemical differentiation, and may result in reduced expression of filoviral cellular receptors. Another possibility is that chemical differentiation drives the expression of a usable filovirus receptor that was not investigated in this study. Given that Zapatero-Belinchón et al used un-differentiated cells, and the current studies employed differentiated SH-SY5Y cells, differentiation might result in the expression of an as-yet-unidentified virus receptor. Further studies, such as complete RNASeq analysis of differentiated vs undifferentiated cells would need to be performed to generate a list of candidates.

Unlike EBOV, MARV grew well in both neuronal cell lines, indicating that whatever restriction EBOV had in BE(2)-M17 cells was not a problem for MARV. One explanation is that MARV infection is possible with lower cellular NPC1 protein levels than EBOV. Evidence to support this is provided by studies by Carette *et al* (2011), who demonstrated that knockout of the NPC1 gene stopped infection with both EBOV and MARV, whilst inhibition of NPC1 through the use of the drug U18666A had a greater effect on EBOV growth than MARV growth, with MARV titres >10-fold higher than EBOV 72h post-infection in the presence of U18666A (55).

Neither EBOV nor MARV appear to induce cell death in infected BE(2)-M17 and SH-SY5Y cells, as evidenced by the cell survival and maintained viral titres beyond Day 10 post-infection. Indeed, the MARV-infected SH-SY5Y cells remained alive with viral titres >10^6^ pfu/ml 21 days post-infection. After 21 days post-infection, the mock-infected SH-SY5Y cells appeared unhealthy, and were similar in appearance to the EBOV-infected cells. By contrast, the MARV-infected SH-SY5Y cells remained alive and maintained their neuronal morphology with interconnecting neurites. Whilst it is possible that MARV is somehow maintaining cell survival of the infected SH-SY5Y cells, it is certainly clear that neither EBOV nor MARV is inducing cell death in either BE(2)-M17 or SH-SY5Y cells the way they do in other cell lines, such as Huh7 cells. Accordingly, it appears that neuropathology during filovirus infection is not a result of direct neuronal death, but instead may result from the response of activated microglial cells interacting with infected neurons. This possibility would explain previous observations of panencephalitis with microglial nodules in human MARV infection (6).

The two non-coding mutations identified in the SH-SY5Y-passaged MARV samples are interesting in that they appear not to be necessary for growth of MARV in SH-SY5Y cells, but were present at consensus level (>90% frequency) in all three replicate passage lineages so must be of benefit to the virus during repeated passaging. The first of the mutations, A2834G, is located at the start of the highly-conserved termination sequence in the gene end of the downstream non-coding region of the NP gene (56, 57). The location of this mutation, in the NP/VP35 gene boundary, and the importance of this region in VP30-mediated transcription in Ebola virus replication, suggests that the mutation may play a role in modulating gene expression downstream of NP (58). The second mutation observed in SH-SY5Y-passaged MARV, G5850A, is in the upstream non-coding region of the GP gene. Given the importance of gene ends in translation efficiency, this mutation might affect the levels of the GP protein within the cell.

Taken together, these results demonstrate that both EBOV and MARV are capable of infecting human neuron-like cells *in vitro*. Moreover, these infections appear to occur in the absence of clear cell death, suggesting the possibility of viral persistence within neurons of the human brain. Viral persistence in neurons has been observed with other viruses, such as measles virus, whereby persistent infection for months to years can lead to subacute sclerosing panencephalitis (SSPE) (59). Given the previously-observed filoviral persistence in other immunoprivileged sites, such as the eye and testes, persistence in neurons *in vivo* appears a real possibility. Further studies into filoviral persistence, including in neurons, will require the development of appropriate animal models that permit the study of long-term sequelae, as well as the development of improved *in vitro* models of human tissues.

## Materials and Methods

### Virus Strains

Zaire ebolavirus (EBOV; Kikwit-95), Reston ebolavirus (RESTV; Pennsylvania-89), and Marburg virus (MARV; Angola-05) were obtained from the NIH Rocky Mountain Laboratory (EBOV and MARV; Hamilton, Montana, USA) or the Centres for Disease Control and Prevention (RESTV; Atlanta, Georgia, USA). Stocks were grown in VeroE6 cells (Sigma Aldrich, Castle Hill, NSW, Australia), with supernatant harvested on Day 7 post-infection. The supernatant was centrifuged at 10,000g for 10min to clarify before being aliquoted into 1ml aliquots. The virus stocks were stored at −80°C until use. All work with infectious virus was performed within the Physical Containment (PC)-4 laboratory at the CSIRO Australian Animal Health Laboratory, Geelong, VIC, Australia.

### Cell Culture and Neuronal Differentiation

VeroE6 and SH-SY5Y cells were obtained from the European Collection of Authenticated Cell Cultures (ECACC; Public Health England, Porton Down, Wiltshire, UK), Huh7 cells were kindly provided by Professor David Jans (Monash University, Clayton, VIC, Australia), and BE(2)-M17 cells were kindly provided by Professor Roberto Cappai (University of Melbourne, Parkville, VIC, Australia).

VeroE6 and Huh7 cells were cultured in Dulbecco’s Modified Eagle Medium (DMEM) GlutaMAX/HEPES medium (Thermo Fisher Scientific, Scoresby, VIC, Australia) containing 8% FCS and 1x Pen/Strep (Thermo Fisher Scientific, Scoresby, VIC, Australia). Following infection, the cells were cultured in the same medium with 2% foetal calf serum (FCS) and 1x Penicillin/Streptomycin.

SH-SY5Y and BE(2)-M17 cells were cultured in neuronal cell Culture Medium: a 1:1 ratio of Minimal Essential Medium (MEM) GlutaMAX and Ham’s F12 GlutaMAX medium (Thermo Fisher Scientific, Scoresby, VIC, Australia) containing 8% FCS and 1× Pen/Strep (Thermo Fisher Scientific, Scoresby, VIC, Australia). Cells were not used beyond five passages post-receipt, as multiple passages can reduce differentiation quality.

For chemical differentiation (based upon Filograna et al, 2015 (37)), 2-3×10^5^ SH-SY5Y or 2-3×10^4^ BE(2)-M17 cells were plated in triplicate 25cm^2^ (T-25) flasks in Culture Medium (defined above; Supplementary Figure 1). After overnight incubation to allow for attachment, the cells were washed with DPBS without Ca^2+^ or Mg^2+^ (Thermo Fisher Scientific, Scoresby, VIC, Australia) and received neuronal cell Differentiation/Infection Medium (1:1 ratio of MEM GlutaMAX and Ham’s F12 GlutaMAX medium containing 2% FCS and 1x Pen/Strep). Flasks of SH-SY5Y cells were treated with 10nM staurosporine in Dimethyl sulphoxide (DMSO; Sigma Aldrich, Castle Hill, NSW, Australia), or the equivalent volume of DMSO only (Sigma Aldrich, Castle Hill, NSW, Australia). Flasks of BE(2)-M17 cells were treated with 10μM *trans*-Retinoic Acid (Sigma Aldrich, Castle Hill, NSW, Australia), or the equivalent volume of DMSO. Differentiation was performed for 7 days, with each flask of cells being washed with DPBS and receiving new medium/drugs on Days 3 and 5 of differentiation.

### RT-qPCR Analysis of Gene Expression

Triplicate flasks of SH-SY5Y and BE(2)-M17 cells were grown in neuronal cell Culture Medium, and were untreated, mock-differentiated (DMSO-only), or drug-treated for differentiation as described above. After the 7-day differentiation, supernatant was removed from each of the flasks, and the cells were lysed by the addition of 1ml TRIzol Reagent (Thermo Fisher Scientific, Scoresby, VIC, Australia). Total RNA was purified from the TRIzol samples using a Zymo Direct-zol™ RNA Mini-Prep kit following the manufacturer’s protocol, including the optional DNase digestion (Zymo Research, Irvine, CA, USA). RNA concentrations in the purified samples were determined using a Qubit RNA HS kit with a Qubit 2 Fluorometer (Thermo Fisher Scientific, Scoresby, VIC, Australia). First-strand DNA synthesis was performed using 200ng purified RNA and SuperScript IV VILO Master Mix with ezDNase Enzyme (Thermo Fisher Scientific, Scoresby, VIC, Australia). qPCR was performed on the cDNA samples using TaqMan Fast Advanced Master Mix (Thermo Fisher Scientific, Scoresby, VIC, Australia) with custom-designed TaqMan Gene Expression Assay plates (Thermo Fisher Scientific, Scoresby, VIC, Australia) using a QuantStudio 6 Flex Real-Time PCR instrument (Thermo Fisher Scientific, Scoresby, VIC, Australia). The plates were designed to allow for the determination of expression of known and possible filovirus receptors, as well as commonly-used markers of human neurons. The gene targets used are listed in Table 2.

**Table 2:**
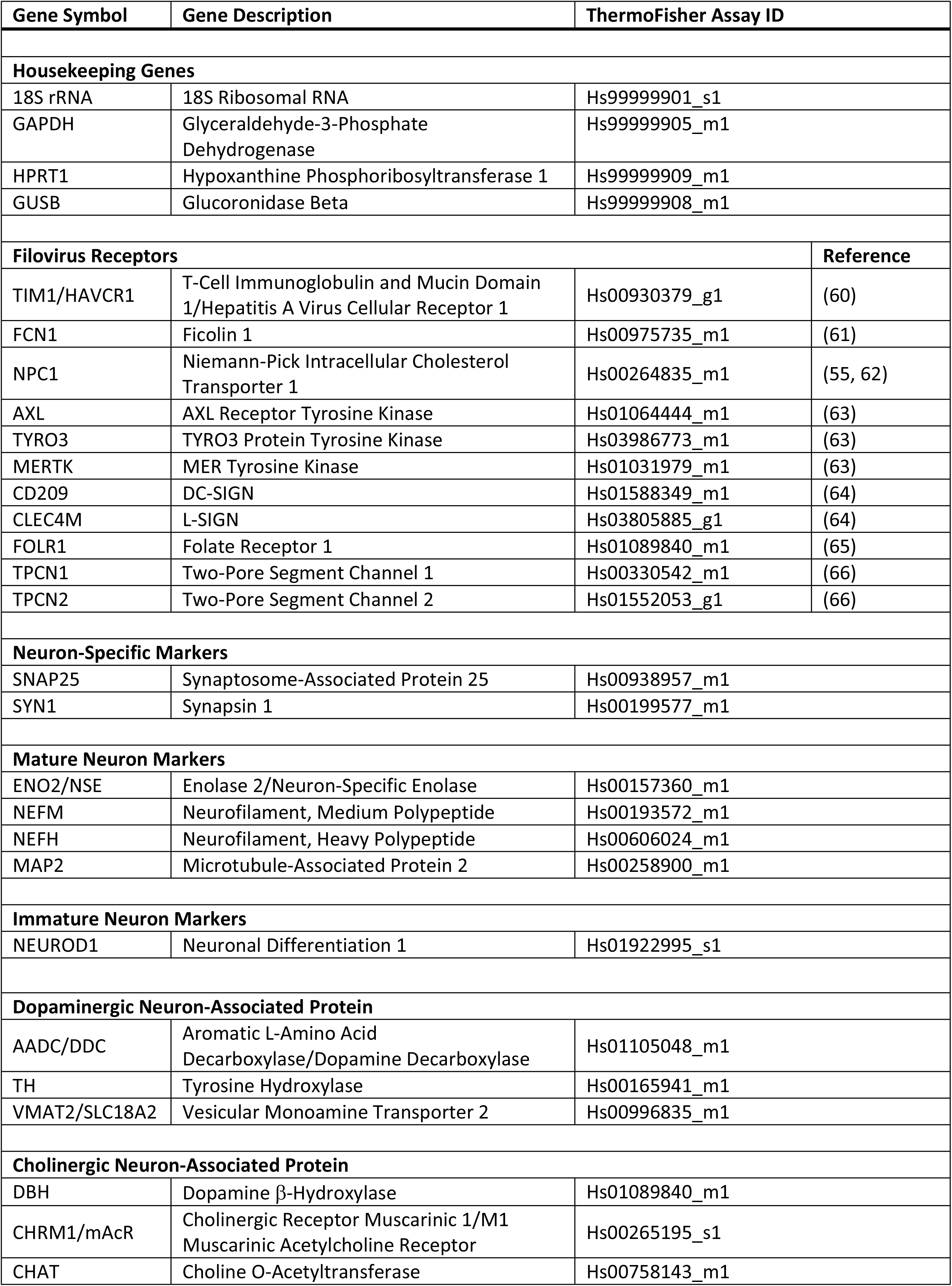
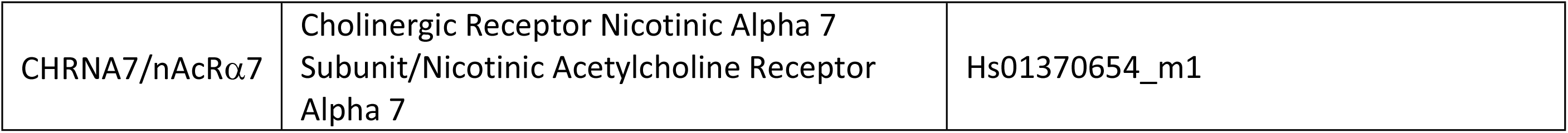
RT-qPCR Targets For Gene Expression Analysis

Ct values obtained from the RT-qPCR arrays were normalised to the geometric mean of the four housekeeping genes. The normalised Ct values were plotted in a heatmap using Mathematica 11 (Wolfram Research, Champaign, IL, USA) scaled between Ct values of 10 (18S RNA-like levels) and 35 (negative cut-off) to generate a semi-quantitative pseudo-northern blot. The normalised Ct values were also used to calculate fold-changes by the ΔΔCt method (67).

### Serial Passaging of Filoviruses in Neuron-Like Cells

Triplicate T-25 flasks of differentiated SH-SY5Y and BE(2)-M17 cells were infected with 1ml Zaire ebolavirus, Reston ebolavirus, or Marburg virus at a multiplicity of infection (MOI) of 0.5. The flasks were incubated at 37°C/5% CO_2_ for 1h with rocking every 10min. At the end of the 1h incubation, cells were washed with DPBS to ensure that any subsequent virus in the supernatant was derived from infected cells. After removal of the PBS, 7ml neuronal cell Differentiation/Infection Medium was added to each flask, and the flasks were incubated at 37°C/5% CO_2_ for 5 days. After incubation, supernatant samples were removed and clarified by centrifugation at 10,000g for 5min. Clarified supernatant samples were stored at −80°C until flasks of cells were ready for the next passage. Serial passaging of the viruses continued for a total of 5 passages, at which point clarified supernatant samples from each passage were titrated.

### Titration of Filovirus Samples

Samples of supernatant from infected cells were titrated by plaque assay (Zaire ebolavirus and Marburg virus) or immunofocus assay (Reston ebolavirus). For both assays, VeroE6 cells were seeded into 6-well plates in Culture Medium 48-96h prior to infection. Cells were seeded at a dilution appropriate to ensure approximately 90% confluence on the day of the assay.

On the day of titration, serial 10-fold dilutions of the virus samples were prepared in DMEM GlutaMAX containing 2%FCS and 1xPen/Strep. Each 6-well plate was washed with PBS, and the serially-diluted samples were added to the appropriate wells (200μl per well). The plates were incubated at 37°C/5% CO_2_ for 30min, with rocking every 10min to ensure that the monolayers did not dry out. At the end of the incubation, 4ml of overlay containing a 1:1 ratio of 2x Eagle’s Minimal Essential Medium (EMEM) with 1x Pen/Strep/Glutamine (Lonza, Mount Waverley, VIC, Australia) and 1.5% agarose (Promega, Alexandria, NSW, Australia) was added to each well. The plates were returned to the incubator, and were incubated at 37°C/5% CO_2_ for 7 days.

For the plaque assays, 2ml of overlay containing 1.5ml Neutral Red solution (Sigma Aldrich, Castle Hill, NSW, Australia) per 50ml overlay was applied to each well at the end of the 7-day incubation. The plates were incubated for a further 24h before plaques were counted and titres (plaque forming units (pfu)/ml) determined.

For the immunofocus assays, the plates were formalin fixed and the agarose plugs removed after the 7-day incubation. The fixed, inactivated plates were removed from the PC-4 facility, and formalin was removed from the wells. The plates were rinsed with water, followed by DPBS with Ca^2+^ and Mg^2+^ (Thermo Fisher Scientific, Scoresby, VIC, Australia). Cells were permeabilised by incubation in 500μl 0.1% NP40 (Sigma Aldrich, Castle Hill, NSW, Australia) in PBS for 30min at room temperature with rocking. The permeabilisation reagent was removed, and the wells were blocked using 0.1% Tropix I-BLOCK (Thermo Fisher, Scoresby, VIC, Australia) in PBS for 1h at room temperature with rocking. The blocking buffer was removed, and the cells were incubated with 400μl of an in-house rabbit anti-Reston Nucleoprotein (NP) antiserum diluted 1:2,000 in blocking buffer. The primary antibody incubation was performed for 1h at room temperature with rocking. The cells were washed 3 times with PBS containing 0.1% Tween-20 (Sigma Aldrich, Castle Hill, NSW, Australia; PBS-T) before being stained with 400μl AlexaFluor 488-labelled anti-Rabbit IgG (Thermo Fisher Scientific, Scoresby, VIC, Australia) at a 1:1,000 concentration in blocking buffer. The plates were incubated at room temperature for 30min with rocking. At the end of the incubation, the plates were washed 3 times with PBS-T, with an additional 2 washes with PBS. Each well received a final 1ml PBS, and the plate was imaged for immunofocus determination using an iBright FL1000 Imager (Thermo Fisher, Scoresby, VIC, Australia). Sample titres were calculated in the same manner as the plaque assay plates.

### Viral RNA Purification and Sequencing

Supernatant samples were inactivated in TRIzol Reagent (Thermo Fisher Scientific, Scoresby, VIC, Australia). Total RNA was purified from the TRIzol samples using a Zymo Direct-zol™ RNA Mini-Prep kit following the manufacturer’s protocol, including the optional DNase digestion (Zymo Research, Irvine, CA, USA). RNA concentrations in the purified samples were determined using a Qubit RNA HS assay kit with a Qubit 2 Fluorometer.

For next-generation sequencing of whole viral genomes, purified viral RNA was concentrated using a RNA Clean & Concentrator™-5 kit (Zymo Research, Irvine, CA, USA) and RNA concentration was determined using a Qubit RNA HS assay kit read on a DeNovix DS-11FX Spectrophotometer/Fluorometer (DeNovix, Wilmington, DE, USA). Total RNA was reverse-transcribed then isothermally amplified using a REPLI-g WTA Single Cell kit (Qiagen, Hilden, Germany) as previously described (68). Dual barcoded Illumina libraries were prepared with the Nextera XT DNA kit according to the manufacturer’s protocol (Illumina, San Diego, CA, USA). Library size distribution was determined using a High-Sensitivity DNA Kit run on a Bioanalyzer 2100 instrument (Agilent Technologies, Santa Clara, CA, USA). Pooled, normalised and denatured libraries spiked with 2% PhiX Control Library were sequenced using the CSIRO AAHL MiniSeq Sequencing System (Illumina, San Diego, CA, USA) using the High-Output Reagent kit (300-cycles), generating 150nt paired-end reads.

### Next-Generation and Sanger Sequencing Data Analysis

*De novo* assembly of complete viral genomes was performed with trimmed, normalised, and error-corrected paired-end reads using the SPAdes genome assembly algorithm provided in Geneious 11 (Biomatters, Auckland, New Zealand). CLC Genomics Workbench 11 (Qiagen, Hilden, Germany) was used to verify *de novo* assemblies by mapping back trimmed paired-end reads with published reference sequences (Zaire ebolavirus: JQ352763; Marburg virus: KY047763). The latter read mapping data was also used to determine variants using default settings of the CLC Basic Variant Detection tool, except that minimum variant detection frequency was reduced to 10%; Ploidy =1; Base Quality Filter =Yes.

For Sanger sequencing, purified RNA was reverse transcribed using AMV Reverse Transcriptase (NEB, Ipswich, MA, USA) with specific primers (IDT, Singapore). Amplicons were generated using Q5 PCR Master Mix (NEB, Ipswich, MA, USA), and the resulting PCR products were cleaned using a NucleoSpin Gel and PCR Clean-Up kit (Macherey-Nagel, Düren, Germany). Samples were prepared for sequencing using a BigDye Terminator 3.1 kit (Thermo Fisher, Scoresby, VIC, Australia), and were sequenced using an Applied Biosystems 3500×l Genetic Analyser (Thermo Fisher Scientific, Scoresby, VIC, Australia). Sequences were aligned to references sequences for analysis using Geneious 11 (Biomatters, Auckland, New Zealand).

### Growth Kinetics in Neuronal and Non-Neuronal Cells

Triplicate T-25 flasks were prepared of Huh7, RA-differentiated BE(2)-M17, and staurosporine-differentiated SH-SY5Y cells. The cells were infected with 1ml stock or Passage 5 Zaire ebolavirus or Marburg virus at an MOI of 0.01. Inocula were back titrated to confirm infectious dose. The flasks were incubated at 37°C/5% CO_2_ for 1h with rocking every 10min. At the end of the 1h incubation, 7ml cell type-appropriate Infection Medium (as outlined above) was added to each flask, and the flasks were rocked to mix. A 500μl aliquot of supernatant was removed from each flask to be the Day 0 samples and was stored at −80°C until titration. Every 24hrs until Day 7, an additional 500μl aliquot of supernatant was removed from each flask and was centrifuged at 10,000g for 5min to clarify before being stored at −80°C until titration. After the removal of the Day 7 sample, the flasks were topped up with 5ml appropriate medium to replace the volume removed during the daily sampling.

By Day 7, the cells in the mock-treated Huh7 flasks were completely dead, and the flasks were discarded with no further samples were taken. For the BE(2)-M17 cells, samples were taken up to, and including, Day 10. Due to the lack of universal cell death in the mock-treated or MARV-infected SH-SY5Y flasks, samples were taken on Days 10, 14, and 21, with a 50% media change on Day 14.

## Acknowledgements

The authors would like to thank Shawn Todd, Matt Bruce, Sarah Edwards, and Jenn Barr for their technical assistance within the PC-4 laboratory.

---

**Supplemental Figure 1:**
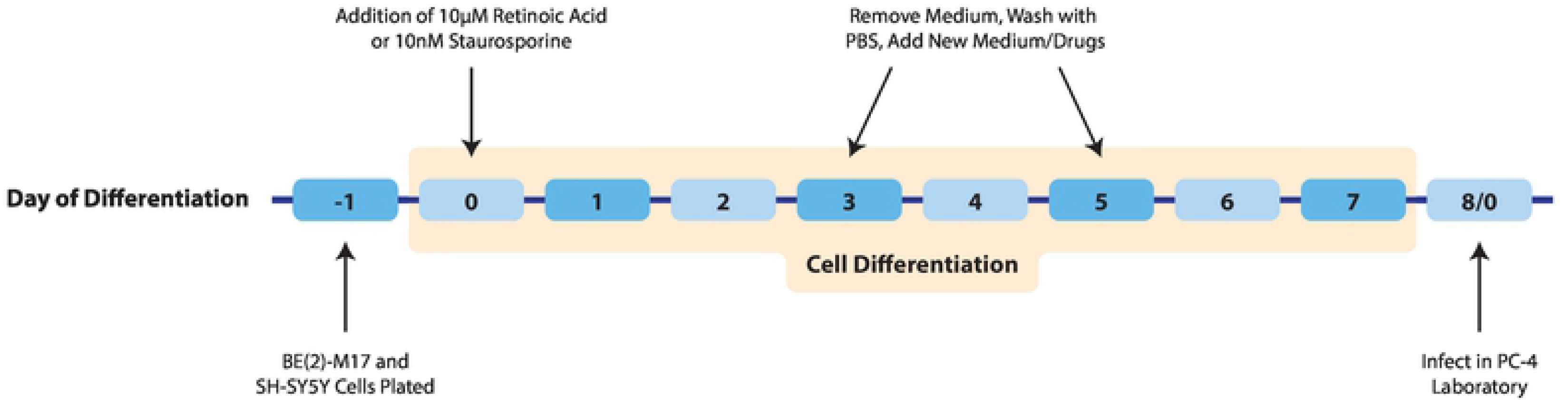
Schematic of BE(2)-M17 and SH-SY5Y Cell Differentiation Protocol

Supplemental Table 1: Mutations Present in Passaged EBOV and MARV Samples (10-35% Frequency)

